# Mutational mimics of allosteric effectors: a genome editing design to validate allosteric drug targets

**DOI:** 10.1101/632232

**Authors:** Qingling Tang, Maria T. Villar, Antonio Artigues, John P. Thyfault, Udayan Apte, Hao Zhu, Kenneth R. Peterson, Aron W. Fenton

**Author notes:** Research in the Fenton, Apte, and Peterson laboratories are funded by NIH grants R01 GM115340, R01 DK098414 and R01 HL111264, respectively. Corresponding author Aron W. Fenton, The University of Kansas Medical Center, Biochemistry and Molecular Biology, MS 3030, 3901 Rainbow Boulevard, Kansas City, Kansas 66160.

## Abstract

Development of drugs that allosterically regulate enzyme functions to treat disease is a costly venture. Screening mutations that mimic allosteric effectors *in vitro* will identify therapeutic regulatory targets enhancing the likelihood of developing a disease treatment at a reasonable cost. We demonstrate the potential of this approach utilizing human liver pyruvate kinase (hLPYK) as a model. Inhibition of hLPYK was the first desired outcome of our screen. We identified individual point mutations that: 1) mimicked allosteric inhibition by alanine, 2) mimicked inhibition by protein phosphorylation, and 3) prevented binding of fructose-1,6-bisphosphate (Fru-1,6-BP). Our second desired screening outcome was activation of hLPYK. We identified individual point mutations that: 1) prevented hLPYK from binding alanine, the allosteric inhibitor, 2) prevented inhibitory protein phosphorylation, or 3) mimicked allosteric activation by Fru-1,6-BP. Combining the three activating point mutations produced a constitutively activated enzyme that was unresponsive to regulators. Expression of a mutant hLPYK transgene containing these three mutations in a mouse model was not lethal. Thus, mutational mimics of allosteric effectors will be useful to confirm whether allosteric activation of hLPYK will control glycolytic flux in the diabetic liver to reduce hepatic glucose production and, in turn, reduce or prevent hyperglycemia.

## Introduction

An emerging class of drugs that allosterically modulate enzymatic activity is generating considerable excitement because of the many reported advantages associated with this approach for the treatment of metabolic diseases ^1–6^. Safety of traditional competitive inhibitor drugs is always an issue, since when present in excessive amounts, they can result in an overdose. Allosteric drugs have an important advantage over competitive inhibitor drugs because once an allosteric drug reaches sufficient concentration to saturate the effector binding site, higher concentrations do not cause additional regulatory effects. Another related advantage is that the extent of the allosteric effect can be moderated based on drug chemistry design ^5,7^. However, despite the promise of the allosteric drug approach, methods to verify that an allosteric response, demonstrated *in vitro*, modifies a disease state *in vivo* are lacking. *In vivo* model systems are needed to verify outcomes of targeting allosteric regulation before committing to the cost of allosteric drug development.

To truly appreciate the need for *in vivo* verification of allosteric drug targets, consider the following scenario. Cell culture or animal studies demonstrate that a given protein is a potential contributor to an observed phenotype. A literature review indicates an allosteric regulation for the identified protein in the context of a signal transduction pathway. Unfortunately, allostery for the isolated protein was characterized at 20°C, pH 8.2, in a very low protein context, and in a hypertonic-Na^+^/buffer solution, an environment which fails to recapitulate *in vivo* physiology. pH, salt concentration, temperature, and other conditions used for these *in vitro* evaluations are usually selected based on the set of parameters that results in the largest detectable allosteric response. Test tube conditions may be quite arbitrary relative to cellular environments, where the extent of allosteric regulation is dependent on pH ^8^, salt type and concentration ^9^, and temperature ^10,11^. The challenges are to accurately incorporate *in vitro* observations to explain *in vivo* outcomes and evaluate test-tube allostery as a relevant regulation that can be modified to treat disease. These *in vitro*/*in vivo* correlation deficiencies are particularly problematic in efforts to rationally design allosteric drugs. Thus, we propose to introduce mutations that mimic allosteric regulation by genome-editing into cell or animal models to verify allosteric drug targets *in vivo*. In this study, we focus on “allosterically engineering” human liver pyruvate kinase (LPYK for liver PYK and hLPYK for the human isozyme) to demonstrate that mutations can mimic allosteric outcomes *in vivo*, the first step needed in our proposed model system design.

Like other PYK isozymes, hLPYK catalyzes the final step of glycolysis, the conversion of phosphoenolpyruvate (PEP) and MgADP to pyruvate and MgATP. The regulatory features of hLPYK embody many examples of classic regulatory mechanisms: 1) Phosphorylation of hLPYK reduces the affinity of this enzyme for the PEP substrate. Thus, protein phosphorylation, catalyzed by cAMP-dependent protein kinase, and dephosphorylation coordinate a balance between inhibition and activation of glycolysis and gluconeogenesis dependent upon cellular energy supply. 2) The fructose-1,6-bisphosphate (Fru-1,6-BP) activation of hLPYK is a classic example of an intermediate from a metabolic pathway acting as a feed-forward activator in the same pathway. 3) Alanine inhibits hLPYK during periods of protein breakdown (*e.g.*, starvation) when the liver uses alanine as the primary carbon source for gluconeogenesis. Regulation alters the affinity of this enzyme for PEP without altering the catalytic rate (*k*_*cat*_) or the affinity for MgADP. In a healthy individual, these regulatory mechanisms result in tight control of PYK activity in the liver. This control is critical since 90% of blood glucose produced by gluconeogenesis is generated in the liver. For the treatment of hyperglycemia, these regulatory mechanisms offer a variety of potential targets for rational drug design aimed at increasing hLPYK activity, either through activation or preventing inhibition, and therefore increasing hepatic glycolysis. More specifically, we anticipate that activation of liver pyruvate kinase under conditions that favor hepatic gluconeogenesis, such as occurs in the diabetic liver, will create a futile cycle that will waste energy and reduce/prevent hepatic production of glucose (Figure 1).

**Figure 1.**
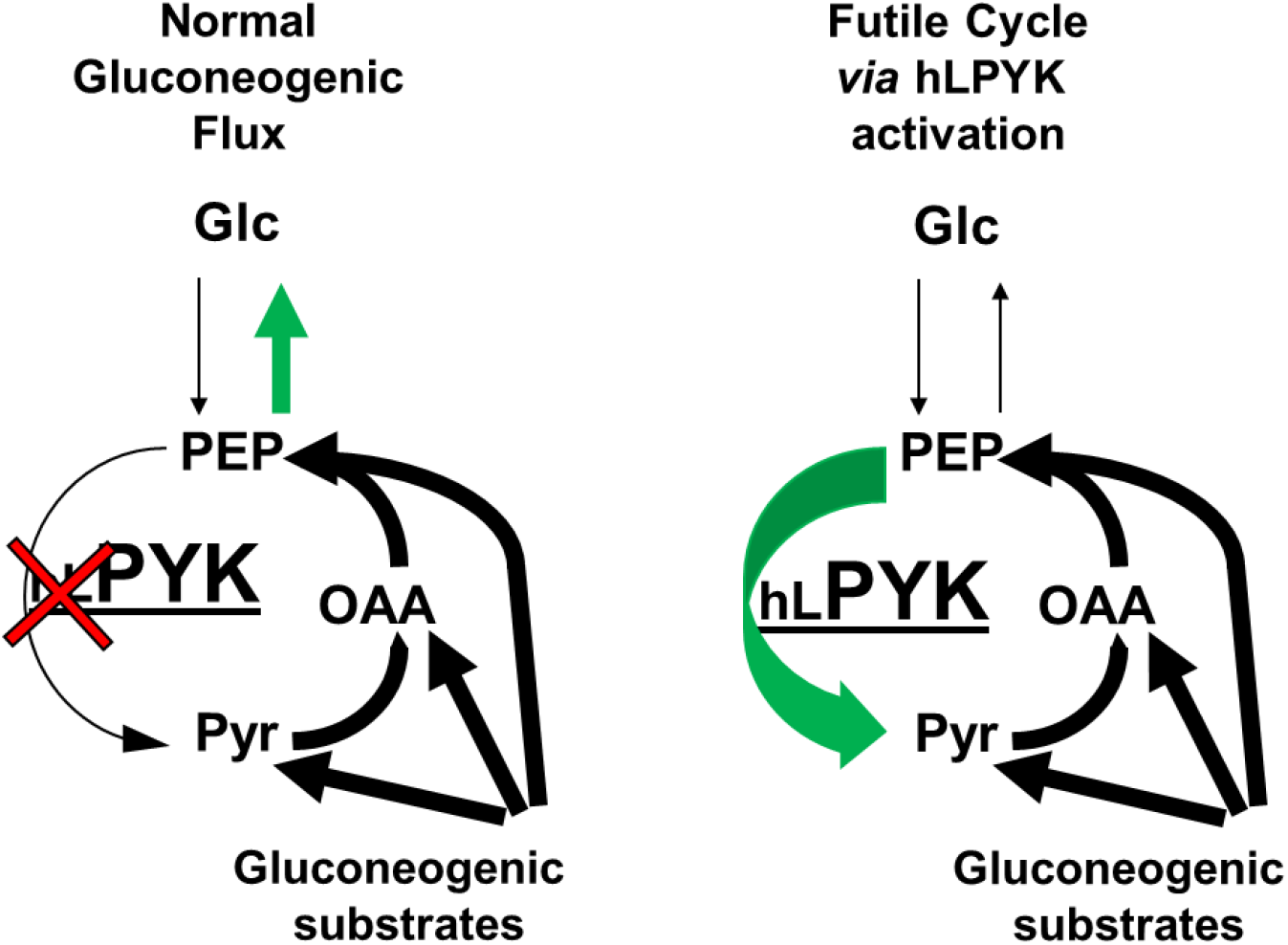
Comparison of expected metabolic flux in the liver with diabetes and elevated gluconeogenesis (left) and in the presence of activated hLPYK (right). When hLPYK is chronically activated, PEP will be converted back to pyruvate (Pyr), “outrunning” gluconeogenic reactions that produce glucose (Glc). Pyruvate is then likely to be used in the TCA cycle for generation of ATP energy, which will, in turn, be used to drive the conversion of pyruvate to oxaloacetate (OAA) and OAA to PEP in an energy wasting futile cycle.

## Materials and Methods

All methods were performed in accordance with relevant guidelines. This includes animal studies, which were performed according to NIH guidelines. The Animal Care & Use Protocol was approved by both KUMC Institutional Biosafety Committee and the KUMC Institutional Animal Care & Use Committee. All use of recombinant DNA according to the NIH Guidelines as approved by the KUMC Institutional Biosafety Committee.

### In vitro characterizations of mutant hLPYK

All protein purification and enzyme assay methods have been previously described ^12^. Mutagenesis of the human *l/r-pyk* gene was performed with a QuikChange kit (Stratagene). Proteins were expressed in the FF50 strain of *Escherichia coli* ^8^ that has both native *E. coli* PYK genes deleted. Mutant proteins were partially purified using ammonium sulfate fractionation followed by dialysis ^13^. Activity measurements were at 30ºC, using a lactate dehydrogenase coupled assay in HEPES buffer, pH 7.5 ^9^. Titrations of activity with a range of concentrations of PEP were used to evaluate *K*_app-PEP_(the PEP concentration at half *V*_*max*_) and that *K*_app-PEP_ value was determined over a concentration range of effector (*i.e.*, Fru-1,6-BP or alanine) to evaluate allosteric coupling.

### Transgene design

A transgene was constructed using standard molecular biology procedures. The development of this mouse model and all studies including mice were approved by the KUMC Institutional Animal Care and Use Committee. All studies were performed according to NIH guidelines and regulations. The *Mus musculus* BAC clone RP23-350K3 was used as a source for gene regulatory elements from the mouse *Pyklr* gene. These elements included 3 kilobases of DNA upstream from the start site (*i.e.*, the *r-pyk* start site, see below) along with the first two exons of this gene. All promoter elements and *cis*-acting DNA elements necessary for transcriptional activation are included in this upstream region ^14–16^. This design, using the native mouse *Pyklr* promoter, assured proper tissue selective gene expression, normal RNA splicing and normal protein expression levels. One caveat of the transgenic approach was that the presence of multiple gene copies or the site of integration of the transgene(s) into the mouse genome was variable and could cause increased protein expression. The beginning of the third exon from the mouse *Pyklr* gene was fused to the cDNA ^17^ encoding the final 10 exons of the human *l/r-pyk* gene. H476L, S531E, and S12A mutations were introduced into the transgene construct as described above. The transgene cassette was isolated for microinjection by restriction enzyme digestion and agarose gel purification.

This transgene cassette was microinjected into the pronuclei of fertilized mouse C57BL/6 oocytes and transgenic mice were obtained, following protocols utilized by the KUMC Transgenic and Gene Targeting Institutional Facility. Germline transmission and transgene lethality were tested by crossing founder male transgenic animals with C57BL/6 females. Due to the constitutively activated design of the transgene, no effort was made to alter the native copy of the mouse *Pyklr* gene. The influence on metabolism we were most interested in is the condition in which wild-type mouse LPYK produced from the native *Pyklr* mouse gene is inhibited, but the hLPYK protein produced from the transgene would remain activated. As noted below, the potential of hybrid LPYK tetramers undermines these considerations and, therefore, the use of the transgenic mouse created in this study was limited to confirming that expression of the triple mutant hLPYK is not lethal.

### Mass spectrometry detection of modified hLPYK expression

Livers from wild-type and transgenic mice were extracted and liver tissue was homogenized with a hand-held ground-glass homogenizer. Samples were filtered through four layers of cheesecloth. Western blot analysis using a polyclonal goat anti-LPYK antibody (a gift from Dr. James Blair) did not reveal any differences in protein expression level.

For multidimensional protein identification (MudPIT), protein samples (~20 mg) were precipitated by adding chilled 100% TCA at 1/10 of the sample volume and incubating overnight at −20°C. Samples were then thawed and centrifuged at 12,000×g at 4°C for 10 min. The pellet was washed with 100 μL of mass spectrometry grade water. 1mL of cold acetone was added to the pellet, and incubated overnight at −20°C. The next day, the pellet was collected by centrifugation at 12,000×g for 10 min at 4°C. The pellets was resuspended in 100 μL of 6 M guanidine hydrochloride in 100 mM ammonium bicarbonate buffer. Protein concentration was measured using a Bradford assay. For each sample, a total 2 mg of protein were reduced and alkylated. Each sample was diluted 6 fold in 100 mM ammonium bicarbonate buffer to reduce the concentration of guanidine hydrochloride to less than 1M. Trypsin was added to 1:10 ratio. Tryptic digestion was allowed to proceed overnight at 37°C. Trypsin digestion was terminated by acidification of the sample with 1% formic acid to below pH 3. All samples were analyzed using MudPIT ^18^. Briefly, peptide digests were loaded on a peptide trap reverse phase C18 column (100 Å, 5 µ, Magic C18 particles, Michrom Bioresources) and desalted from the peptide trap for 5 min with buffer A (0.1% formic acid in HPLC-grade water). Peptides were eluted from the peptide trap using a 40-min gradient of 0–40% buffer B (0.1% formic acid in HPLC-grade acetonitrile) followed by a 10-min gradient of 40–98% buffer, and loaded on line onto the SCX part of a 100 μm ID biphasic column packed in house with 3 cm of polysulfoethyl A strong cation exchange (SCX) matrix (5 µm, 300 Å, Poly LC) and 12 cm of reverse-phase resin (100 Å, 5 µ, Magic C18 particles, Michrom Bioresources). The peptides were step-eluted from the SCX phase onto the reverse phase of the biphasic column using 2-min salt pulses of 10, 20, 30, 40, 50, 60, 70, 80, 90, 90, 100, and 100% of buffer C (500 mM ammonium acetate, 5% acetonitrile, and 0.1% formic acid), respectively. For the second dimension, each fraction from the SCX chromatographic step, the peptides were separated with a 0 – 60 % (v/v) acetonitrile gradient in 0.1 % (v/v) formic acid in 90 minutes at a flow rate of 0.25 µl/min. The mass spectrometer (LTQ FT, ThermoFisher Scientific) was operated in positive ionization mode with a proxeon nanospray source operated at 2.5 Kv and source temperature of 250°C. The instrument was operated in data-dependent acquisition mode to perform survey mass scan analysis on the FT followed by MSMS scans on the ion trap of the ten most intense ions, with a dynamic exclusion of two repeat scans of the same ion, 30 sec repeat duration and 90 sec exclusion duration. Fragment ion spectra were acquired using normalized collision energy of 35% and a maximum injection time of 100 ms. For data analysis, all MSMS scans were searched using Peaks Studio 8.5 (BSI) or Proteome discoverer (version 2.0, Thermo Fisher Scientific) running the Sequest HT algorithm. A database search was conducted against a *Mus musculus* protein database derived from the NIBInr repository as of January 9, 2017, to which the mutant for of the human pyruvate kinase was added. The refined data were subjected to database search using trypsin cleavage specificity, with a maximum of 2 missed cleavages. The following variable modifications were selected: pyroglutamination from Q and E (N-terminal), oxidation of M, and deamidation of N, Q. Carboxymethylation of C was selected as a fixed modification, and a maximum of 3 modifications/peptide was allowed. Estimation of false positive rate (FDR) was conducted by searching all spectra against a decoy database consisting of the inverted sequences of all proteins in the original (direct) database. For peptide identification a FDR ≤ 1 was defined and a minimum of two unique peptides per protein was required for protein identification. Amino acid sequence assignment of all peptides of interest was subsequently inspected manually. A total of 17,910 peptide sequences were identified, corresponding to 1,715 protein groups.

## Results

The site of phosphorylation and locations of ligand binding sites have been mapped onto the structure of PYK ^5,17,19–26^. In previous studies, we used knowledge of these locations to introduce a series of mutations into each regulatory site ^12,13,27^. Those mutation series empirically identified mutations that both increase and decrease allosteric responses (Figure 1). However, for use in *in vivo* studies for drug development, it will be important for mutations to influence only the targeted allosteric functions without causing global changes to all enzyme functions. Therefore, in this study we extended the characterization of previously identified mutations to ensure that each mutation influences only a single regulatory property of the enzyme. We also extended the study by combining three mutations that promote more activity for both an *in vitro* characterization of protein regulation and an *in vivo* test to confirm that expression of the constitutively active enzyme is not lethal in mice.

H476L, S531E, and S12A mutations all convert the enzyme towards a more activated/less inhibited form that is, the enzyme binds PEP with a higher affinity ^12,13,27^. H476L causes a complete loss of response to the allosteric inhibitor alanine. Based on previous data ^7^, this lack of response is likely due to the inability of the H476L mutant protein to bind alanine or any other regulatory amino acid. However, without direct information on alanine binding to the protein, a lack of response to an effector does not distinguish whether the effector fails to bind to the mutant protein or if the effector binds, but does not elicit an allosteric response. The H476L mutation in the amino acid effector binding site does not influence PEP affinity in the absence of effectors (left y-intercept in graphs of Figure 2) or modify the response of the enzyme to Fru-1,6-BP activation.

**Figure 2.**
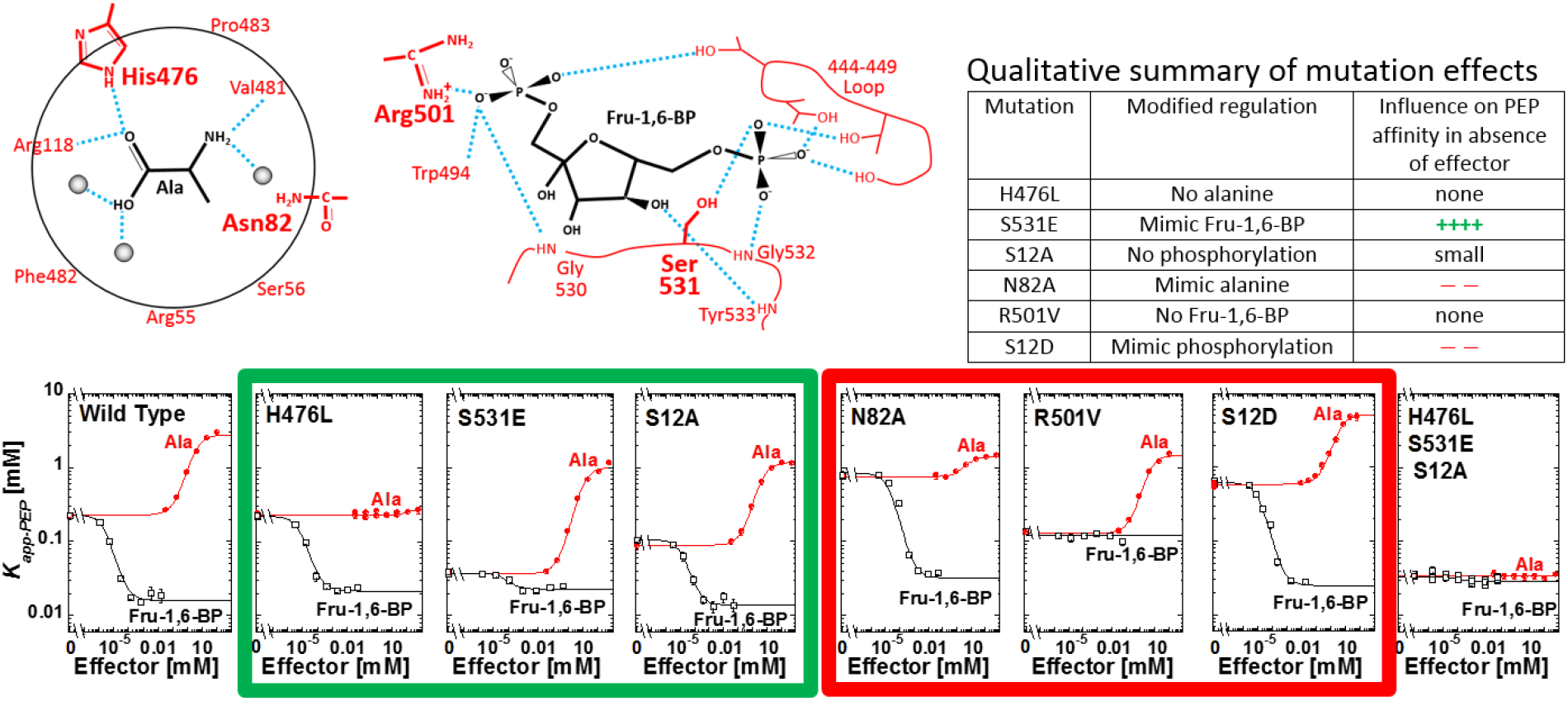
Top: Schematic of effector binding sites for alanine (left) and Fru-1,6-BP (center). Locations of mutations used in this study are in bold. A qualitative summary of the effects of each mutant is included at top right. Bottom: Allosteric effects of alanine (●;red) and Fru-1,6-BP (□;black) on the PEP affinity (*K*_*app-PEP*_) on wild-type and mutant hLPYK enzymes. The amino acid binding site was defined using rabbit M1-PYK^5^ however, residues are mostly conserved, thus, this figure includes hLPYK residue labels. Mutations originally identified and partially characterized in: H476L ^7^; N82A ^27^; S531E and R501V ^12^; and S12A ^20^. Characterization of S12D was previously reported ^13^.

S531E adds a negative charge to a loop within the Fru-1,6-BP binding site that is critical for allosteric activation. The added charge largely mimics activation by Fru-1,6-BP; that is, PEP affinity in the mutant protein is increased to close to the same affinity as is observed for the wild-type hLPYK protein when Fru-1,6-BP is bound. Furthermore, the addition of Fru-1,6-BP caused only a slight additional increase in PEP affinity. The S531E does not completely activate the enzyme to the same level as Fru-1,6-BP. However, our laboratory has now characterized more than 800 mutations of *l/r-pyk* ^12,27,28^. Using these mutations as a point-of-comparison, the response of S531E is a good mimic of Fru-1,6-BP activation. The S531E mutant protein also retains the response to alanine, although that response (Figure 2, the distance between the plateaus at low and high alanine concentrations) is slightly increased compared to the wild-type enzyme.

The S12A mutation removes the side-chain hydroxyl group of Ser12 that is phosphorylated by cyclic-AMP dependent protein kinase in response to hormone signals ^13^. Although this mutation also slightly reduces PEP affinity (left y-intercept of Figure 2) compared to the-wild-type enzyme, increased PEP affinity influences the enzyme in the same direction as our global goal of a more active enzyme. The S12A enzyme retains qualitative responses to both alanine and Fru-1,6-BP. Quantitatively, the allosteric response (Figure 2, the distance between the plateaus at low and the high effector concentration) to alanine is similar to that of the wild-type enzyme. The allosteric response to Fru-1,6-BP is reduced compared to the wild-type protein due to the slightly reduced PEP affinity in the absence of effector. Nonetheless, the affinity of S12A for PEP in the presence of saturating concentrations of Fru-1,6-BP is similar to the wild-type protein.

Although our main interest is activation of hLPYK, there may be reasons to allosterically engineer this protein for inhibition. N82A, R501V, and S12D are three mutations in the respective three effector sites that will be useful should inhibition become a future focus of our study ^12,13,27^. N82A resides in the alanine binding site. This mutation causes reduced PEP affinity even in the absence of effector (left y-intercept of Figure 2). Addition of alanine resulted in only a modest increase of inhibition. R501V causes a complete loss of response to the allosteric inhibitor alanine. Based on previous data ^12^, this lack of response is likely due to the inability of the R501V mutant protein to bind Fru-1,6-BP. However, as noted earlier, a lack of response to an effector does not distinguish between whether the effector fails to bind to the mutant protein or if the effector binds, but does not elicit an allosteric response. We previously reported that S12D mimics the quantitatively small, yet well documented, inhibition by protein phosphorylation ^13^. Finally, we previously described *in vitro* evidence for protein oxidation of hLPYK that occurs on Cys436^20^. Should this oxidation be confirmed as a regulatory mechanism *in vivo*, then the C436M and C436S mutations will serve to either remove the oxidation, or mimic the oxidation, respectively. These mutations are available for allosterically engineering other regulatory motifs into the enzyme as needed in future studies.

Our long-term goal is to determine if activation of hLPYK is a valid drug target for activating hepatic glycolysis and reducing hepatic glucose production (gluconeogenesis) as a treatment for the hyperglycemia associated with diabetes. Genome-editing the mutations that mimic activation by Fru-1,6-BP (S531E), that prevent inhibition by alanine (H476L), and that prevent inhibition by phosphorylation (S12A) should result in a fully activated enzyme to facilitate the *in vivo* evaluation of the control of LPYK regulation on hyperglycemia. As anticipated, the triple mutant enzyme binds PEP, even in the absence of effector, similar to the Fru-1,6-BP activated wild-type enzyme. In addition, this modified protein does not respond to either alanine or Fru-1,6-BP. Thus, the triple mutant protein exemplifies an “allosterically engineered” protein that may be useful in an *in vivo* model to validate hLPYK as an anti-hyperglycemic drug target.

We chose to first test lethality of this fully activated form of the enzyme by expressing a *l/r-pyk* construct that included the H476L, S531E and S12A mutations in a transgenic mouse line as outlined in Figure 3. Due to the sequence and size similarity between mouse and human LPYK, western blot analysis did not confirm expression of the transgene protein product. However, these analyses indicated that similar total LPYK protein expression levels were observed in transgenic and wild-type C57BL/6 mice, that is, presence of the transgene did not increase the protein expression level (data not shown). To demonstrate that the transgene protein product was being expressed, we completed a MudPIT Mass spectrometry analysis of liver samples from transgenic mice (Figure 4). However, this analysis was challenging given the high sequence similarity between human and mouse LPYK. The protein coded for in the transgene was identified in transgenic mice, but not control mice. The MudPIT evaluation did not show whether the protein products from the transgene and the native *pyklr* mouse gene are expressed at the same level.

**Figure 3.**
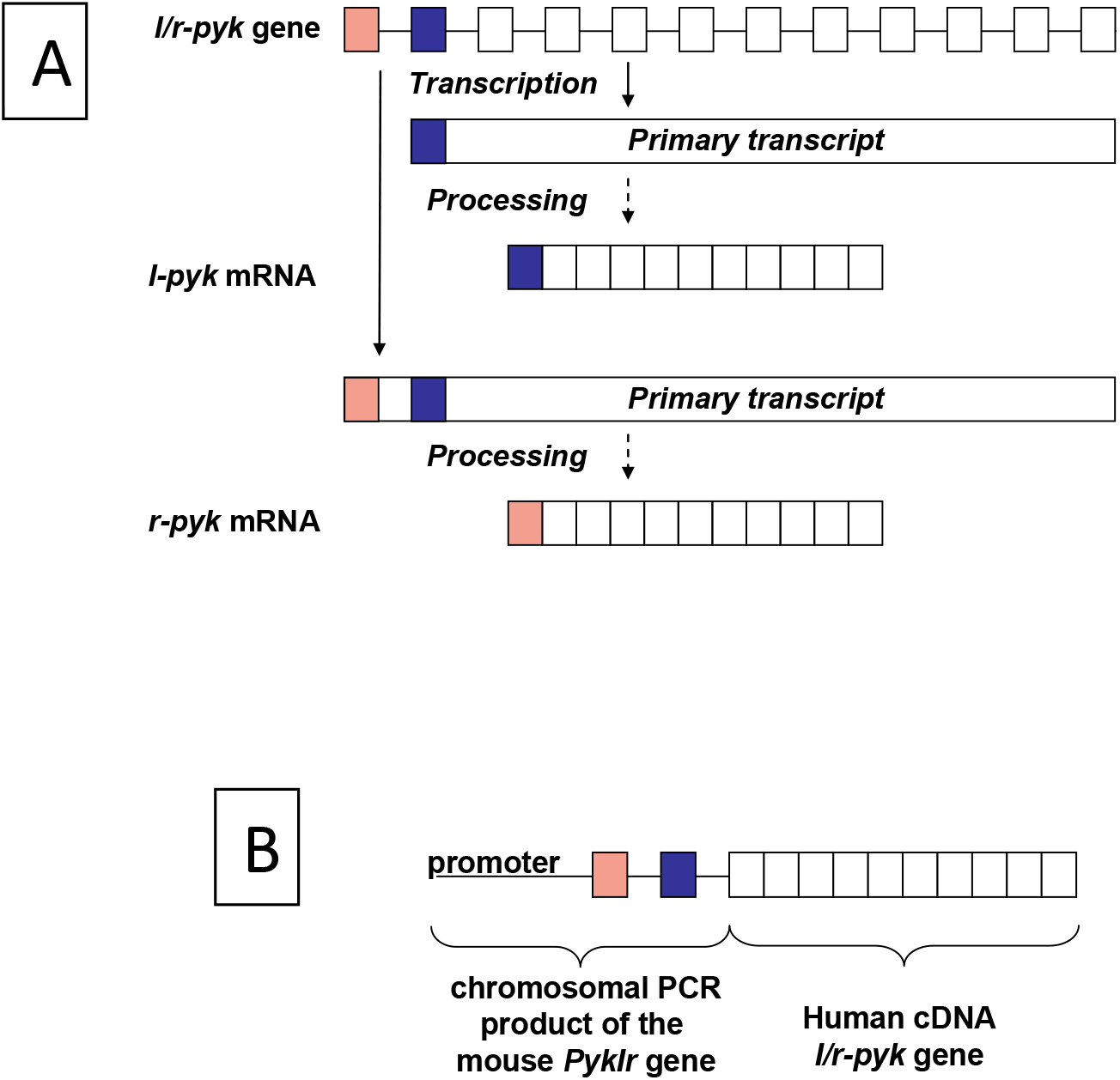
Schematic of the *l/r-pyk* expression transgene construct. A) The *l/r-pyk* gene gives rise to two gene product (R-PYK expressed in erythrocytes and L-PYK expressed in liver) due to the use of different start sites. The three mutations H476L, S531E, and S12A were all included in the human cDNA portion of the transgene. B) The design of the transgene used in this study to construct a new transgenic mouse line.

**Figure 4.**
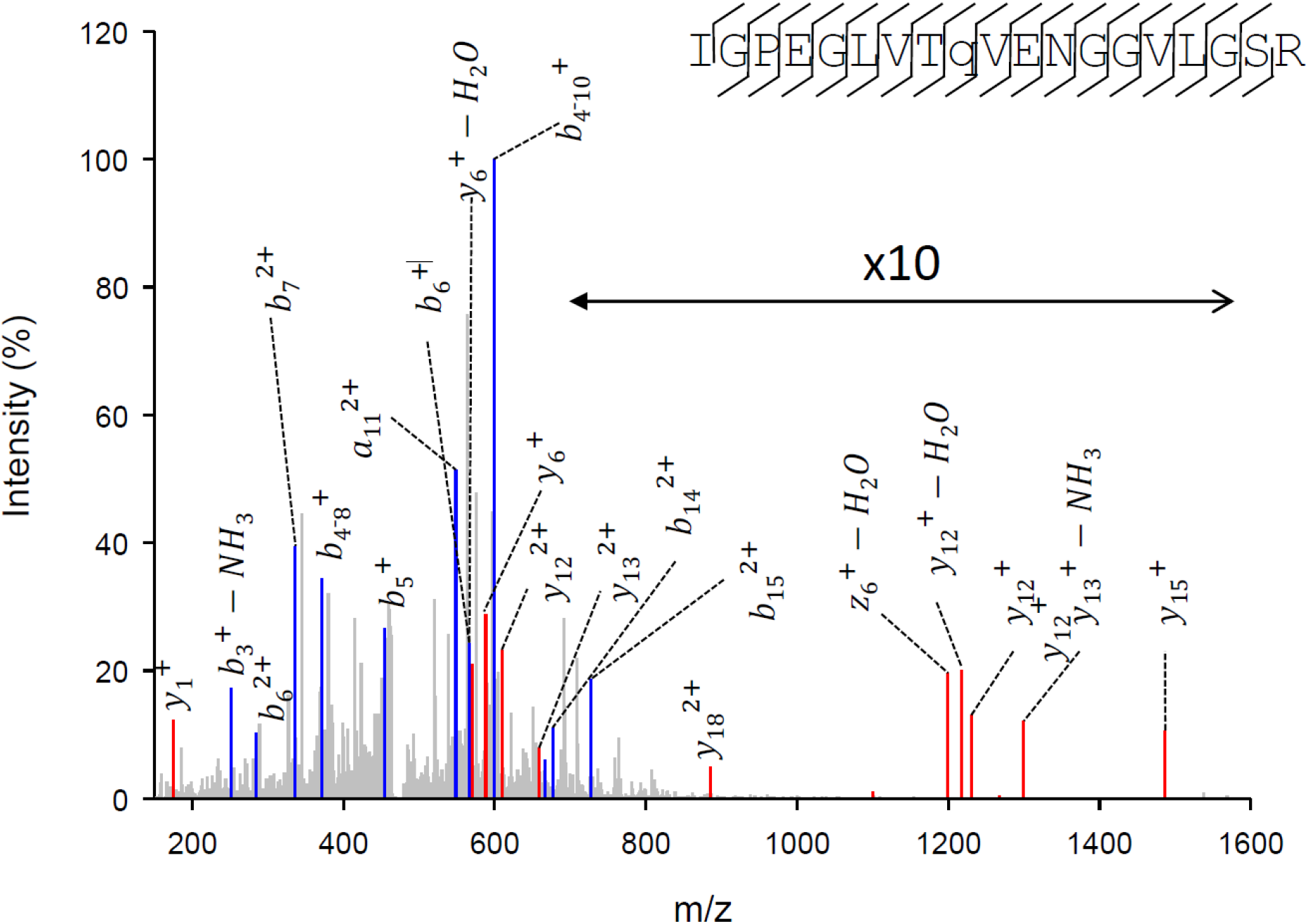
Identification of the human liver pyruvate kinase protein expressed in the liver of the transgenic mouse. The CID tandem mass spectrum of the quadruple-charged ion with m/z of 471.5029. This is the only unique peptide identified for human LPYK. For clarity, not all the fragment ions observed are indicated. Amino terminal and carboxyl-terminal ions are highlighted in blue and red, respectively. The fragmentation pattern is indicated on the corresponding peptide sequence (upper right). This peptide was identified by Peaks as indicated in Material and Methods, at a −10logP of 23.4, corresponding at a FDR of 0.7 %, and 1.6 ppm mass error.

Base on the survival of the transgenic mice, we concluded that expression of the constitutively activated hLPYK subunits was not lethal *in vivo*. Importantly, the test for lethality was the only reason we created the transgenic mouse line. LPYK is a homotetramer and human and mouse LPYK protein sequences are highly similar. In our design, the triple mutant human LPYK protein is expressed in the presence of expression of the mouse LPYK protein. The fully activated hLPYK protein product from the transgene should be dominant to the wild-type mouse LPYK, which can be inhibited. However, the potential hybrid tetramers between wild-type mouse LPYK subunits and the mutated human LPYK subunits may mask the total activation associated with the human subunits. Therefore, further studies were not pursued with this mouse model. The fact that mutant transgene expression was not lethal provides a way forward for future transgenic and/or genome-edit models to study LPYK function in mouse models.

## Discussion

The long-standing tools of the geneticist to knock-out, knock-down and/or overexpress a protein are not useful to evaluate allostery *in vivo*. Novel methods are needed to verify outcomes of targeting allosteric regulation *in vivo*. We propose that knocking-in point mutations that mimic allosteric effectors in mice using CRISPR/Cas9 genome-editing may partially fill this need. Genome-editing will only partially fill the need since constitutive modification of allosteric properties is likely to elicit a compensatory change in gene expression. Therefore, the constitutive nature of genome-edited mutations will not completely mimic the acute response to allosteric effectors. The genome-editing approach we propose requires detailed knowledge about the outcomes of individual mutations on allosteric functions. Although our understanding of allosteric mechanisms lags our understanding of other protein structure/function questions, we demonstrate that we can empirically identify mutations that mimic that results from binding an allosteric effector. Thus, we anticipate that knocking-in mutations in cell culture and animal models will be useful to model potential outcomes from allosteric drugs.

All mutations included in the current work are in one of the regulatory sites of *l/r-pyk*. Importantly, each mutation largely modifies only a single type of regulation of the protein. In addition, for mutational mimics, the magnitude of the response (*i.e.*, the extent of change in *K*_app-PEP_) is like those caused by the respective native allosteric effectors that interact with the wild-type protein. We speculate that these mutations (Figure 2) modify PEP affinity in the active site by preventing or mimicking the respective native allosteric/covalent mechanism. Our H476L/S531E/S12A triple mutant protein data provide evidence that the individual mutations, each with a specific influence on allosteric properties, can be combined to maximize this effect. This enzyme binds PEP similar to the Fru-1,6-BP activated wild-type protein. Furthermore, it does not respond to either alanine or Fru-1,6-BP addition. Therefore, the triply mutated enzyme is maximally activated and is not further regulated.

Directly transitioning from in vitro protein characterization to developing a mouse model without evaluating the influence of modifications in cell culture is particularly applicable in the study of hLPYK. Even when derived from hepatocarcinomas, cell cultures in general fail to express hLPYK.

The transgenic mouse model created with the triple mutant *l/r-pyk* gene thrives; expression of the mutant protein does not cause lethality. Live extracts from those mice confirm expression of a modified protein from the transgene. Therefore, the S531E, H476L, and S12A mutations, or combinations of these will be very useful to understand which type of hLPYK regulation is to be targeted for drug design to treat hyperglycemia. These outcomes of this exploratory study support the feasibility of future studies to evaluate if activation of glycolysis in the diabetic liver will be useful to prevent/reduce hepatic glucose production for the treatment of hyperglycemia.

## Author contribution

Qingling Tang: Ms. Tang made mutant genes and expressed proteins for in vitro characterization.

Maria T. Villar: Dr. Villar worked with Dr. Artigues in proteomics evaluations of protein expression.

Antonio Artigues: Dr. Artigues worked with Dr. Villar in proteomics evaluations of protein expression.

John P. Thyfault: Drs. Thyfault, Apte and Zhu worked together to develop the transgenic mouse model.

Udayan Apte: Drs. Thyfault, Apte and Zhu worked together to develop the transgenic mouse model.

Hao Zhu: Drs. Thyfault, Apte and Zhu worked together to develop the transgenic mouse model.

Kenneth R. Peterson: Dr. Peterson and Dr. Fenton designed the project and directed all aspects of the project.

Aron W. Fenton: Dr. Fenton designed the project, directed all aspects of the project, and contributed directly to enzymatic assays.

## Data availability statement

All data included in this manuscript will be made available upon request.

## Relevant guidelines

The mouse model generated in this work was done with the approval and under the supervision of the Institutional Animal Care and Use Committee (IACUC) at The University of Kansas Medical Center.

## Acknowledgments

Funding in the Fenton, Apte, and Peterson laboratories are funded by NIH grants R01 GM115340, R01 DK098414 and R01 HL111264, respectively. This study used services from both the KUMC Transgenic and Gene Targeting Institutional Facility (directed by Drs. Jay Vivian and Melissa Larson) and the KUMC Mass Spectrometry/Proteomics Core Laboratory (directed by Dr. Antonio Artigues).

